# Simple and rapid site-specific integration of multiple heterologous DNAs into the *Escherichia coli* chromosome

**DOI:** 10.1101/2022.09.13.507873

**Authors:** Lauren A. Riley, Irenee C. Payne, Melissa Tumen-Velasquez, Adam M. Guss

## Abstract

*Escherichia coli* is the most studied and well understood microorganism, but research in this system can still be limited by available genetic tools, including the ability to rapidly integrate multiple DNA constructs efficiently into the chromosome. Site-specific, large serine recombinases can be useful tools, catalyzing a single, unidirectional recombination event between two specific DNA sequences, *attB* and *attP*, without requiring host proteins for functionality. Using these recombinases, we have developed a system to integrate up to twelve genetic constructs sequentially and stably into in the *E. coli* chromosome. A cassette of *attB* sites was inserted into the chromosome and the corresponding recombinases were cloned onto temperature sensitive plasmids to mediate recombination between a non-replicating, *attP-* containing “cargo” plasmid and the corresponding *attB* site on the chromosome. The efficiency of DNA insertion into the *E. coli* chromosome was approximately 10^7^ cfu/μg DNA for six of the recombinases when the competent cells already contained the recombinase-expressing plasmid and approximately 10^5^ cfu/μg DNA or higher when the recombinase-expressing plasmid and “cargo” plasmid were co-transformed. The “cargo” plasmid contains ΦC31 recombination sites flanking the antibiotic gene, allowing for resistance markers to be removed and reused following transient expression of the ΦC31 recombinase. As an example of the utility of this system, eight DNA methyltransferases from *Clostridium clariflavum* 4-2a were inserted into the *E. coli* chromosome to methylate plasmid DNA for evasion of the *C. clariflavum* restriction systems, enabling the first demonstration of transformation of this cellulose-degrading species.

**Importance:** More rapid genetic tools can help accelerate strain engineering, even in advanced hosts like *Escherichia coli*. Here, we adapt a suite of site-specific recombinases to enable simple, rapid, and highly efficient site-specific integration of heterologous DNA into the chromosome. This utility of this system was demonstrated by sequential insertion of eight DNA methyltransferases into the *E. coli* chromosome, allowing plasmid DNA to be protected from restriction in *Clostridium clariflavum* and enabling genetic transformation of this organism. This integration system should also be highly portable into non-model organisms.

## INTRODUCTION

*Escherichia coli* is one of the most important model organisms for bioengineering, and it is arguably the most well-understood and studied [1, 2]. A large genetic toolbox has been developed for *E. coli*, including methods for insertion of heterologous DNA into the chromosome. These tools include homologous recombination, λ Red recombinase-based recombineering and its newer CRISPR/Cas variants [3–5], and site-specific recombinases for heterologous DNA integration into the chromosome [6, 7]. Despite the genetic tools already available, research in *E. coli* can still be hindered by the speed and simplicity of genome engineering tools available. Homologous recombination tools to insert plasmid DNA into the chromosome typically take several steps and selections. Due to the labor and time involved, this technique is not often used in *E. coli*. The λ Red recombinase system and CRISPR/Cas systems greatly improve upon homologous recombination in *E. coli* and allows for the creation of markerless mutants. However, integration efficiency is limited for these systems for DNA fragments larger than 3 kb [3].

Site-specific recombinase systems such as the Conditional-Replication, Integration, and Modular plasmid (CRIM) system represent an efficient method for inserting DNA into the chromosome. They use site-specific DNA recombinases, in this case from native *E. coli* bacteriophages such as λ, to integrate plasmids into specific sites in the chromosome [8]. However, this system does not provide a way to remove the plasmid backbone and the selectable marker, and it is limited to five native phage insertion sites.

Site-specific recombinases catalyze a single recombination event between two specific DNA sequences, which enables DNA integration and excision events to occur [9]. They are classified into two families based on the catalytic amino acid in the active site, tyrosine- or serine-recombinases. The tyrosine family includes the commonly used Cre and Flp recombinases and the bacteriophage λ integrase (λ Int). Cre and Flp recognize identical recombination sites (*loxP* and *frt*, respectively), and therefore recombination is reversible [10]. λ Int catalyzes a single crossover event at two non-identical sites, *attB* (<u>atta</u>chment site from the <u>B</u>acterium) and *attP* (<u>atta</u>chment site from the <u>P</u>hage), to create two new sites, *attL* and *attR* (<u>L</u>eft and <u>R</u>ight), and so recombination is irreversible and stable [11]. The *att* sites from this type of recombinase are about 350-450 bp in size with a common 15 bp core where recombination occurs [11, 12]. λ Int also requires accessory protein IHF (Integration Host Factor) for functionality, and therefore only functions in *E. coli* and very close relatives [12].

The second family of recombinases includes serine recombinases, such as the archetypical large serine recombinase ΦC31 integrase (ΦC31 Int) [7, 9, 11]. These serine recombinases offer benefits over recombinases like Cre, Flp, and λ Int. They function similarly to the λ Int by enabling unidirectional recombination at *attB* and *attP* sites to create *attL* and *attR* sites. However, the *att* sites from these recombinases are typically shorter, ranging in size from 35-100 bp [11]. The unidirectionality enables a stable integration event that will not spontaneously excise from the genome. Furthermore, unlike λ Int, this group does not require any host-specific factors. Therefore, large serine recombinases from phylogenetically diverse sources function in a wide variety of heterologous hosts, and they have been developed as genetic tools in diverse members of the Bacteria, Archaea, and Eukaryota [6, 7, 13–15]. They have also been used in *E. coli* as synthetic biology “parts” [16, 17]. Beyond the ΦC31 Int, there are several well characterized serine recombinases that have been shown to function outside of their native host [7]. These characteristics allow large serine recombinases to be especially useful in genome engineering.

Some applications require more than five successive insertions of heterologous DNA into the *E. coli* chromosome, exceeding the capacity of the CRIM system. One such application is the heterologous expression of DNA methyltransferases to enable the transformation of new bacteria. Restriction-modification (RM) systems have been demonstrated to be an important barrier to bacterial transformation because they recognize and degrade foreign DNA [18–23]. Most RM systems are comprised of a restriction enzyme that targets and hydrolyzes non-methylated DNA at a specific DNA motif and a corresponding methyltransferase that protects the host genome by methylating DNA at the same motif [24]. Bacteria often encode more than one RM system, and to successfully transform an organism, these RM systems need to be evaded through proper methylation of DNA prior to transformation [22]. An example organism is *Clostridium clariflavum* strain 4-2a, an organism capable of degrading lignocellulosic biomass via the production of an enzyme complex called the cellulosome and catabolizing both glucose and xylose at thermophilic temperatures [25]. This strain has not yet been transformed, which is likely due to the presence of eight predicted RM systems encoded in the genome. Thus, evasion of the *C. clariflavum* RM systems likely requires the expression of more than five DNA methyltransferases in *E. coli*.

We sought to expand upon the CRIM system by leveraging large serine recombinases to develop a system for efficiently inserting heterologous DNA into the *E. coli* chromosome. By designing the system to allow for simple removal of the selectable marker and vector backbone, our system enables successive integration of up to twelve constructs without the accumulation of repetitive DNA sequences and selectable markers. As a demonstration of this system in *E. coli*, we expressed eight methyltransferases encoded in the *C. clariflavum* 4-2a genome methylate DNA and enable transformation *of C. clariflavum* for the first time.

## MATERIALS AND METHODS

### Strains and Culture Conditions

*E. coli* Top 10 *Δdcm::frt* was used for maintenance of all replicating vectors and integration of the *attB* cassette [21]. PIR-dependent plasmids, containing the oriR6K origin of replication, were propagated in *E. coli* GT115 (Invivogen). All *E. coli* strains were grown in Luria-Bertani (LB) broth with antibiotics, as necessary, for maintenance of vectors and for selection of integrants. Carbenicillin was used at 50 μg/mL, kanamycin was used at 50 μg/mL for replicating vectors and 30 μg/mL for single copy integrants, and spectinomycin and streptomycin were used together at 100 μg/mL each for replicating vectors and 50 μg/mL each for single copy integrants. All strains were grown at 37°C, unless the strain contains a temperature sensitive recombinase helper plasmid, in which case growth was at 30°C. *Clostridium clariflavum* 4-2a was grown in a Coy anaerobic chamber (Coy Laboratory Products, Grass Lake, MI) in CTFUD medium [21] at 50°C. CTFUD medium was supplemented with 5 μg/mL thiamphenicol when needed.

### Plasmid Construction

Annotated plasmid maps for all plasmids used in this study are provided in the Supplemental Material. All plasmids and strains used in this study are listed in Supplemental Table S1. The first set of recombinase helper plasmids were constructed by GenScript, where the gene encoding λ Int in pINT-ts [8] was replaced with the synthesized serine recombinases that were previously described [7], resulting in plasmids pGSs037, 038, 040, 041, 042, 043, 044, 045, 046, 047, 048, 050, 053, 054, and 082 (Supplemental Table S1). Recombinases were codon optimized for expression in *E. coli*.

For the second set of helper plasmids, recombinases were cloned under the *Clostridium thermocellum* DSM 1313 enolase promoter [26] with a ribosome binding site modified to AGGAGGA. First, ΦC31 was synthesized with the enolase promoter into a pUC replicating vector. The promoter and recombinase DNA were then amplified by polymerase chain reaction (PCR) using Phusion High Fidelity master mix (Thermo Scientific) and inserted using Gibson Assembly (New England Biolabs) into pINT-ts, replacing the native promoter and lambda recombinase, resulting in pLAR047. Using pLAR047 as a backbone, the remaining serine recombinases were cloned and inserted, replacing ΦC31, using Gibson Assembly, resulting in plasmids pLAR052-62 and 074 (Supplemental Table S1).

The poly-*attB* cassette was synthesized by GenScript with all fourteen attachment sites previously described [7] (*att* sites are listed in Supplemental Table S2) except ΦC31 (plasmid pAMGs177). In order, the *attB* sites are Φ370, BT1, R4, BxB1, TP901-1, RV, SpβC, TG1, ΦC1, MR11, ΦK38, A118, Wβ, and BL3. The *attB* cassette was PCR amplified and inserted into the multiple cloning site of pAH144 [8] using Gibson Assembly, resulting in plasmid pLAR080. The *attB* cassette was then integrated using the CRIM system into the HK022 phage *attB* site [8] of 1) *E. coli* Top 10 *Δdcm::frt*, 2) BW25113 *ΔmcrA::frt ΔmcrC-mrr::frt Δdcm::frt*, and 3) BW25113 *ΔmcrA::frt ΔmcrC-mrr::frt Δdcm::frt Δdam::frt*, creating strains AG2005, AG3525, and AG4277 respectively.

The base integration plasmid, pGSs009, was synthesized and constructed by GenScript. It was constructed from pAH55 [8] and a synthesized insert containing ΦC31 *attB* and *attP* sites flanking the T5lac promoter and lacZα. The promoter in pGSs009 was replaced with P_BAD_ from pLA2 [8] to create pLAR067. For recombinase efficiency experiments, a synthesized cassette containing 13 *attP* sites, excluding SpβC, was first inserted into the BamHI and NdeI sites of pGSs009 using Gibson Assembly. Next, oligonucleotides with the Spβ *attP* and homology to pGSs009 were inserted into the EcoO1091 site, creating pLAR031.

*C. clariflavum* methyltransferases (CloclaDRAFT _2368_2369, CloclaDRAFT _1830_1831, CloclaDRAFT _1994, CloclaDRAFT _1058, CloclaDRAFT _1051, CloclaDRAFT _1868_1869, CloclaDRAFT _2133, and CloclaDRAFT _1961) were codon optimized and synthesized by GenScript, each with a different *attP* site. Each methyltransferase was then cloned into pLAR067, under the control of P_BAD_, using NEB builder, resulting in plasmids pMTV37-44 (Supplemental Table S1).

### Testing Integration Efficiencies in Two-Steps

To test integration efficiency of the first set of recombinase helper plasmids, each plasmid was transformed into the poly-*attB* cassette strain, AG2005. Batches of electrocompetent cells were made in duplicate for each strain, and 200 ng of pLAR031 was electroporated into 25 μl of each competent cell batch, each in duplicate (n=4). Cells were resuspended in 250 μl SOC and recovered at 37°C for 1 hour and 42°C for 30 minutes. The recovery was plated on LB with kanamycin and incubated at 37°C overnight. Colonies were picked into 5 mL LB with kanamycin and PCR screened after growth. PCR was performed on strains using primers P1-4 (all screening primers are listed in Supplemental Table S3) to screen for the corresponding recombination event using Quick-Load *taq* master mix (NEB).

### Testing Integration Efficiencies in One-Step

To test integration efficiency of co-transformation of the putative constitutively expressed recombinase helper plasmids and cargo plasmid, batches of electrocompetent cells were prepared of the poly-*attB* cassette strain (*E. coli* strain AG2005) in duplicate. Both the helper plasmid and pLAR031 were co-transformed in duplicate for each batch of competent cells (n=4), using 200 ng of each plasmid and 25 μl of electrocompetent cells. Transformants were resuspended in 250 μl of SOC medium and recovered at 37°C for 1 hour. The recovery was plated on LB + kanamycin plates and incubated at 37°C overnight.

### Removal of Plasmid Backbone

To remove the plasmid backbone, electrocompetent cells were made from the confirmed integration strain and transformed with the ΦC31 helper plasmid. Cells were recovered at 30°C for 1 hour and plated on LB + carbenicillin plates. Colonies were picked into LB (with no antibiotics) and grown at 37°C for 8 hours to allow plasmid curing. Then, a 5 μl aliquot of the outgrowth was streaked on LB plates and grown overnight. Resulting colonies were then patched on LB, LB + carbenicillin, and LB + kanamycin plates to screen for loss of the kanamycin resistance marker and loss of the helper plasmid.

### Methyltransferase expression and methylome analysis

Methyltransferase expression plasmids were sequentially integrated into *E. coli* strain AG4277 as described above, ultimately yielding strain AG5645. Strain AG5645 was grown with 1 mM arabinose to induce methyltransferase expression. Genomic DNA was isolated from *C. clariflavum* and AG5645 using the QIAGEN Genomic-tip kit (QIAGEN). Methylome analysis was performed on these strains using Pacific Biosciences (PacBio) Single Molecule Real Time (SMRT) sequencing [27] and Whole Genome Bisulfite Sequencing (WGBS) as previously described [21].

### Transformation of C. clariflavum 4-2a

Electroporation was performed similarly to existing protocols for *Clostridium thermocellum [28]*. Two 5 mL cultures were inoculated with *C. clariflavum* 4-2a and grown overnight. The next day, two 200 mL cultures were inoculated with a 1% inoculum and grown until an optical density at 600 nm (OD_600_) of 0.9. Once the cultures were grown, they were centrifuged in 50 ml conical tubes at room temperature at 6,000 x g for 15 minutes. The supernatant was decanted, and 25 ml of electroporation buffer (250 mM sucrose, 10% glycerol) was added to the tube without disrupting the cell pellet. Cells were centrifuged again and washed twice more in the same way. After the last spin, the cell pellets were resuspended in ~100 μl electroporation buffer and transferred to a microcentrifuge tube.

Using fresh electrocompetent cells, 20 μl of cells were transformed with 1 μg of pMTL83151 [29]. Cells were electroporated in a 1 mm cuvette with a square wave using a Bio-Rad Gene Pulser Xcell Electroporation System set at 1200 V with a 1.5 msec pulse. Cells were then resuspended in 1 ml CTFUD medium and incubated for 3 hours at 50°C to recover. After recovery, transformants were plated within CTFUD with 1.5% agar and 5 μg/ml thiamphenicol and incubated for 4 days at 50°C. Colonies from the plates were picked into liquid CTFUD medium. Liquid culture was screened by PCR for the presence of the *cat* gene and further confirmed via 16S rRNA gene sequencing to verify the culture. Two batches of *C. clariflavum* competent cells were transformed with plasmid from each methylation state, each time in duplicate (n=4 total).

## RESULTS

### Design of the integration system

A heterologous DNA integration system using serine recombinases was created by first integrating a cluster of *attB* sites into the *E. coli* chromosome at the HK022 phage site using the CRIM system (Figure 1). The resulting strain contains 14 orthogonal *attB* sites stably inserted into the chromosome: Φ370, ΦBT1, R4, BxB1, TP901-1, RV, Spβc, TG1, ΦC1, MR11, ΦK38, A118, Wβ, and BL3.

**Figure 1:**
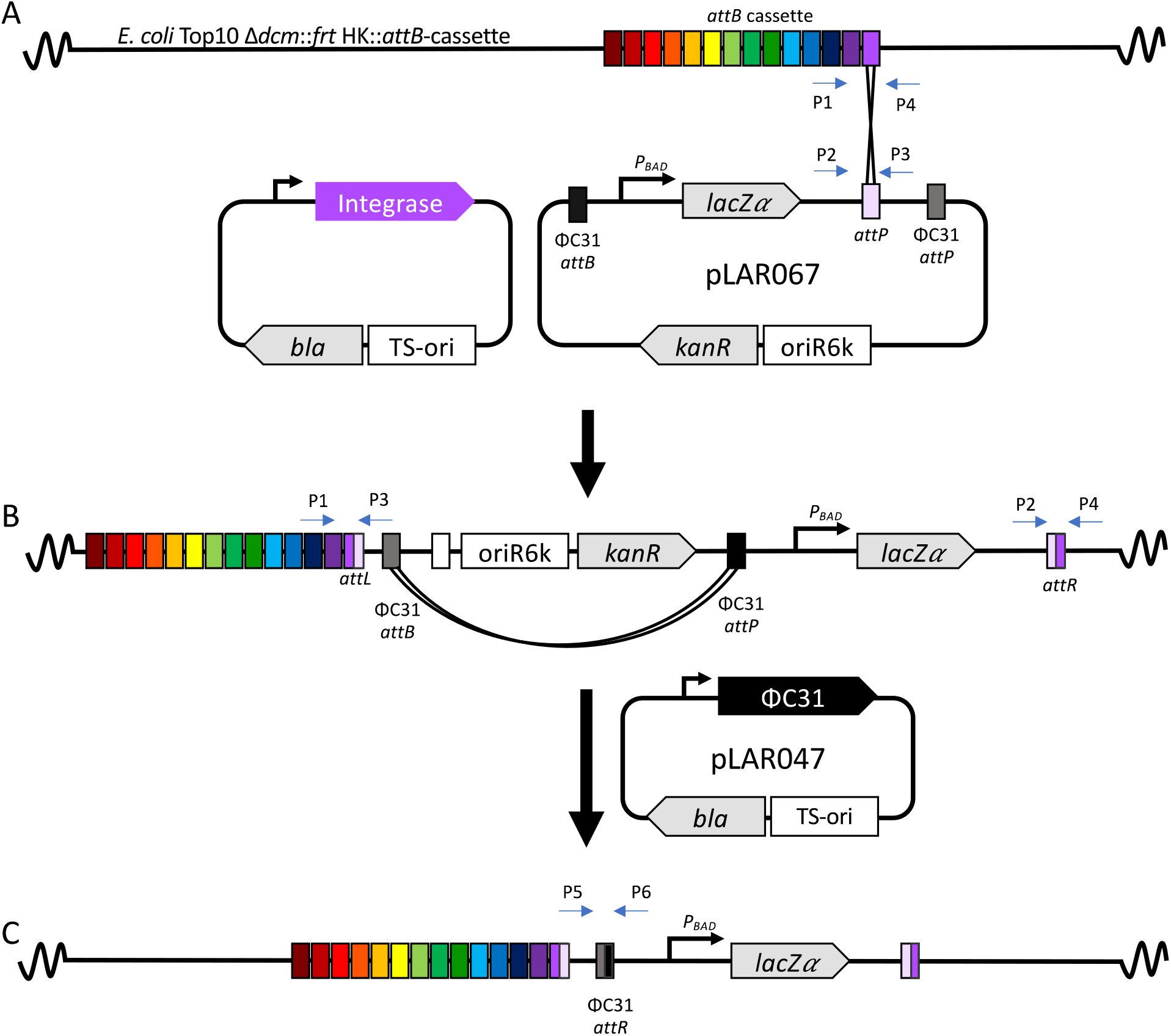
Overview of the site-specific recombinase system for DNA integration into the *E. coli* chromosome. A) 14 *attB* sites were added to chromosome at the HK phage site (AG2005, AG3525, and AG4277) and the 14 corresponding serine recombinases were cloned onto temperature sensitive plasmids to mediate integration between a plasmid containing an *attP* site (e.g., pLAR067) and the corresponding *attB* site on the chromosome. B) The kanamycin resistance marker and plasmid origin of replication can then be removed by introducing the ΦC31 recombinase on a temperature sensitive plasmid (pLAR047).

A non-replicating integration vector, which carries the “cargo” DNA to be inserted, contains the kanamycin resistance marker, the conditional oriR6K origin of replication that only replicates in an *E. coli* strain that expresses the *pir* gene, and one of the corresponding *attP* sites. The *oriR6K-kanR* cassette is flanked by ΦC31 *attB* and *attP* sites, which facilitates marker removal (see below). Outside of the ΦC31 sites is an arabinose-inducible promoter driving lacZα in plasmid pLAR067. This serves as a convenient site for cloning of the intended cargo, with NdeI and BamHI sites flanking lacZα.

Each of the corresponding large serine recombinases were cloned into a temperature-sensitive helper plasmid, pInt-Ts [8], replacing the λ integrase. These genes are under the control of the λ pR promoter and cI857, a heat-inducible promoter system. This enables transient expression of the recombinase to mediate integration and, after raising the temperature, subsequent loss of the recombinase plasmid. The recombinase helper plasmid mediates DNA recombination between an *attP* site from the non-replicating “cargo” plasmid and its associated *attB* on the *E. coli* chromosome, resulting in the integration of the entire non-replicating plasmid into the chromosome. Recombination creates two new attachment sites called *attL* and *attR* that flank the newly integrated DNA. (Figure 1a). Next, the *oriR6K-kanR* cassette can be removed by transient expression of the ΦC31 integrase. This results in recombination between the ΦC31 *attB* and *attP* sites, leaving behind a single ΦC31 *attR* site and any associated genetic cargo (Figure 1B-C). To consistently test each recombinase, a single integration plasmid was constructed that contains each of the corresponding *attP* sites (poly-*attP* cassette; pLAR31).

### Testing Integration Efficiency of the Recombinase helper Vectors

To test integration efficiency of the recombinases, the temperature sensitive helper plasmids encoding recombinases were each transformed into an *E. coli* strain that contains the poly-*attB* cassette on the chromosome. Next, a non-replicating plasmid containing the poly-*attP* cassette (pLAR031) was transformed into each strain expressing a recombinase (Fig. 2). Integration was observed with twelve of the recombinases, of which six (A118, TG1, BL3, Wβ, TP901, and Spβ) had an efficiency of approximately 10^7^ CFU/μg of DNA. Another four showed recombination frequencies in the range of 10^5^ – 10^6^ CFU/μg of DNA. ΦC1 enabled the lowest recombination frequency of an average of 200 CFU/μg of DNA, while two recombinases, ΦBT and RV, did not enable any integration.

**Figure 2:**
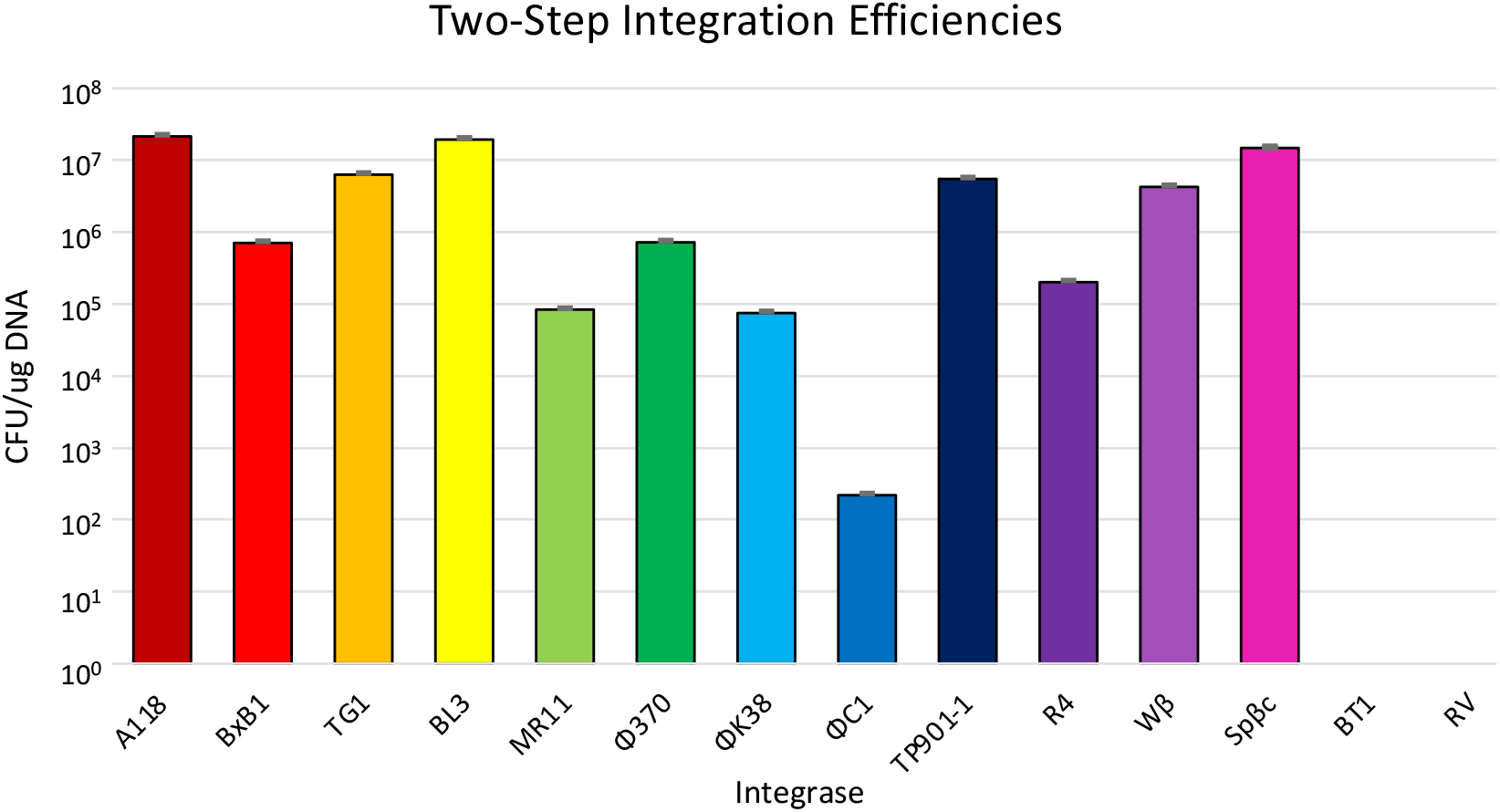
Integration efficiencies of each recombinase in colony forming units per μg (CFU/μg) of the integrating *attP* vector. Error bars represent the standard deviation of four biological replicates.

### Integration Confirmation

Serine recombinases have been shown to integrate DNA into naturally occurring secondary *attB* sites, called pseudo-*attB* sites, that have similarity to the native sequences [30]. To determine if the non-replicating plasmid integrated into the intended locus, rather than a pseudo-*att* site, colonies from the resulting transformations were screened using a four-primer multiplexed PCR screen, with primers flanking the two *att* sites (P1, P2, P3, P4 indicated in Figure 1A), similar to the screen used for the CRIM system [8]. The four-primer system gives distinct band sizes to distinguish between the *attB* (P1 and P4), *attP* (P2 and P3), *attL* (P1 and P3), and *attR* (P2 and P4) sites to evaluate if integration occurred at the intended locus. Upon correct insertion, bands consistent with the *attL* and *attR* will be observed. However, if the poly-*attP* cassette plasmid integrated into a pseudo-*attB* site, a single band, indicative of the parent strain, will form. Ten colonies from each transformation were screened and all colonies showed the correct integration bands for all 10 colonies except MR11 and ΦC1, where 7 of 10 and 4 of 10 integrated at the correct locus, respectively (Supplemental Figure 1).

### Streamlined integration system via co-transformation

To determine if the process could be further streamlined to accelerate strain construction, we explored whether co-transformation of the recombinase helper plasmid and the integrating plasmid could allow for DNA insertion into the chromosome. In this scenario, the recombinase is transiently expressed, but without selection for the helper plasmid. If successful, this would eliminate one round of competent cell preparation, transformation, and plasmid curing. When using the temperature-sensitive helper plasmids described above in combination with integrating poly-*attP* cassette plasmid at a non-permissive temperature, no kanamycin-resistant colonies were observed, indicating no recombination. Expression of the recombinases could have been too low to enable DNA integration; therefore, the temperature-sensitive recombinase helper plasmids were reconstructed to use the *Clostridium thermocellum* enolase promoter (P*_Ct_eno_*), which has been shown to be highly active in *E. coli* [26] and is presumably constitutively expressed. Of the twelve recombinases shown to be functional in Figure 2, the promoter was successfully replaced in all except Spβc and Wβ, where the cloning was not successful.

To test the efficiency of these new recombinase expression plasmids, each were co-transformed with the poly-*attP* cassette vector into the *E. coli* poly-*attB* cassette strain and recovered at a temperature too high to allow replication of the helper plasmid. Each of the 10 enabled integration in the one-step integration process (Fig. 3). Five of the recombinases had an insertional efficiency of 10^5^ CFU/μg of DNA or higher: BxB1, TG1, BL3, Φ370, and R4. The ΦC1 and TP901 recombinases also enabled DNA integration in a single step but had low efficiencies of less than 100 CFU/μg of DNA.

**Figure 3:**
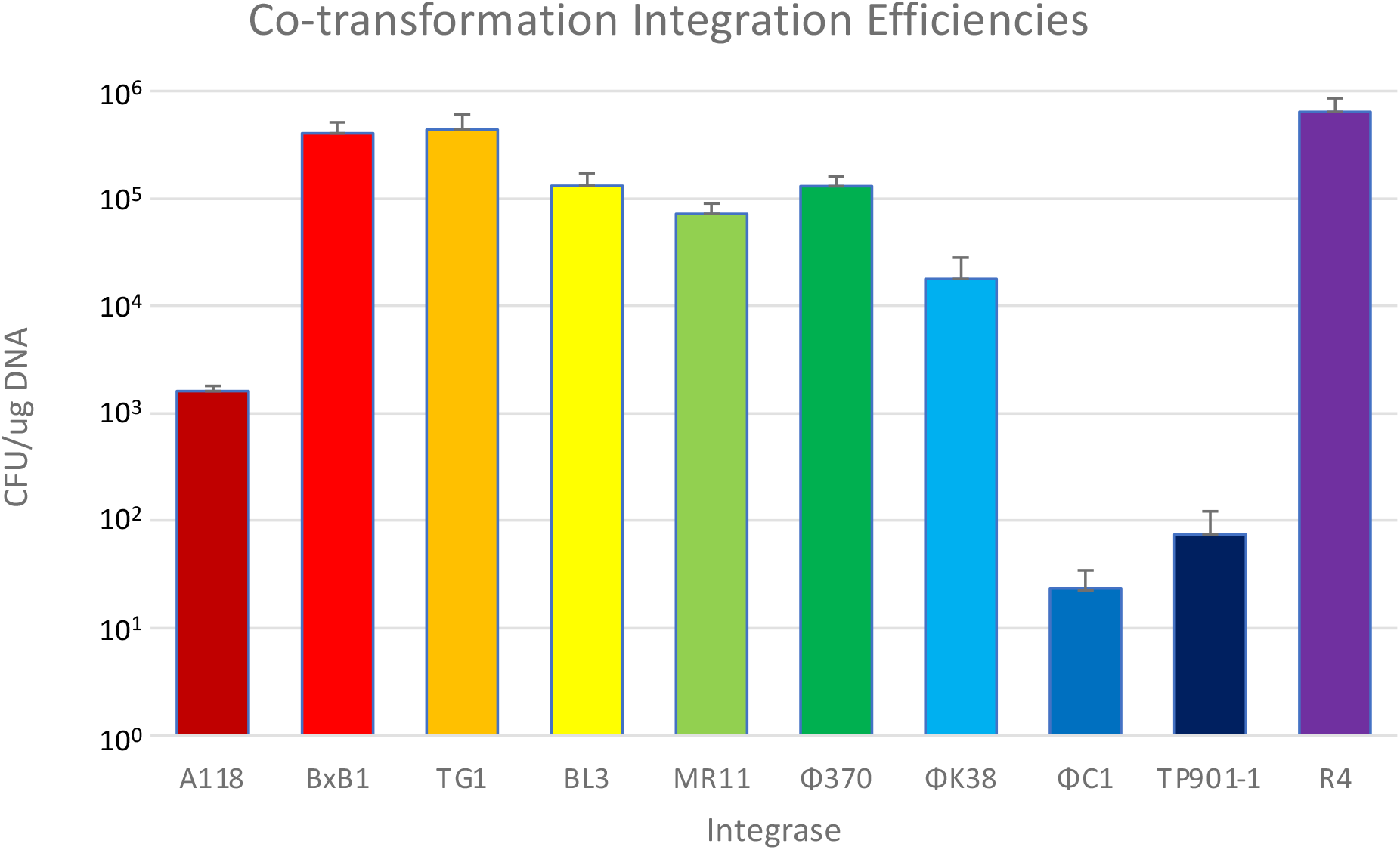
Recombination efficiencies of each second-generation recombinase in colony forming units per μg of suicide *attP* vector (CFU/μg of DNA). Error bars represent the standard deviation of four biological replicates.

### oriR6K-kanR Backbone Removal

Some applications will require the removal of the antibiotic resistance gene, for instance to enable further genetic modification. Therefore, we tested the ΦC31 recombinase-based marker-removal system described above using a strain in which the *attP*-cassette plasmid integrated into the R4 *attB* locus (Figure 1B,C). After integration into the R4 *attB* site was confirmed, a temperature-sensitive plasmid encoding ΦC31 Int was transformed into the strain at 30°C. After further incubation at a 37°C to the cure the ΦC31 helper plasmid, 12 colonies were screened for successful recombination between the ΦC31 *att* sites and loss of the helper plasmid by patching single colonies on each antibiotic used in the process (kanamycin and carbenicillin). Ten colonies (83%) did not grow on either antibiotic and were PCR screened for removal of the backbone using primers P5 and P6 (Figure 1B) that flank the ΦC31 sites. Of the colonies that were sensitive to both antibiotics, 100% were confirmed to have lost the *oriR6K-kanR* cassette by PCR (Supplemental figure 1).

### Identification and expression of methyltransferases for C. clariflavum 4-2a

A major barrier to the genetic transformation of non-model microorganisms is the presence of restriction-modification (RM) systems that degrade foreign DNA. One approach to evading these RM systems is to express the target organism’s DNA methyltransferases in *E. coli*. Therefore, as a demonstration of the utility of the above integration system, we targeted heterologous expression of *C. clariflavum* DNA methyltransferases to methylate plasmids prior to transformation.

Putative methyltransferases in *C. clariflavum* 4-2a were first identified using the NEB Restriction Enzyme Database (REBASE) [31] and genome annotations (GenBank Accession number CP003065.1; [32]). Eight DNA methyltransferase gene clusters were identified: CloclaDRAFT_2368 - 2369, CloclaDRAFT _1830 - 1831, CloclaDRAFT _1994, CloclaDRAFT _1058, CloclaDRAFT 1051, CloclaDRAFT _1868 - 1869, CloclaDRAFT _2133, and CloclaDRAFT _1961. To determine which methyltransferases are functional, methylome analysis was performed using PacBio SMRT analysis to identify motifs containing 6-methyladenine (m^6^A) and 4-methylcytosine (m^4^C) and Whole Genome Bisulfite Sequencing (WGBS) to identify 5-methylcytosine (m^5^C). Methylome analysis revealed eight m^6^A motifs and no m^4^C or m^5^C motifs (Table 1).

**Table 1:**
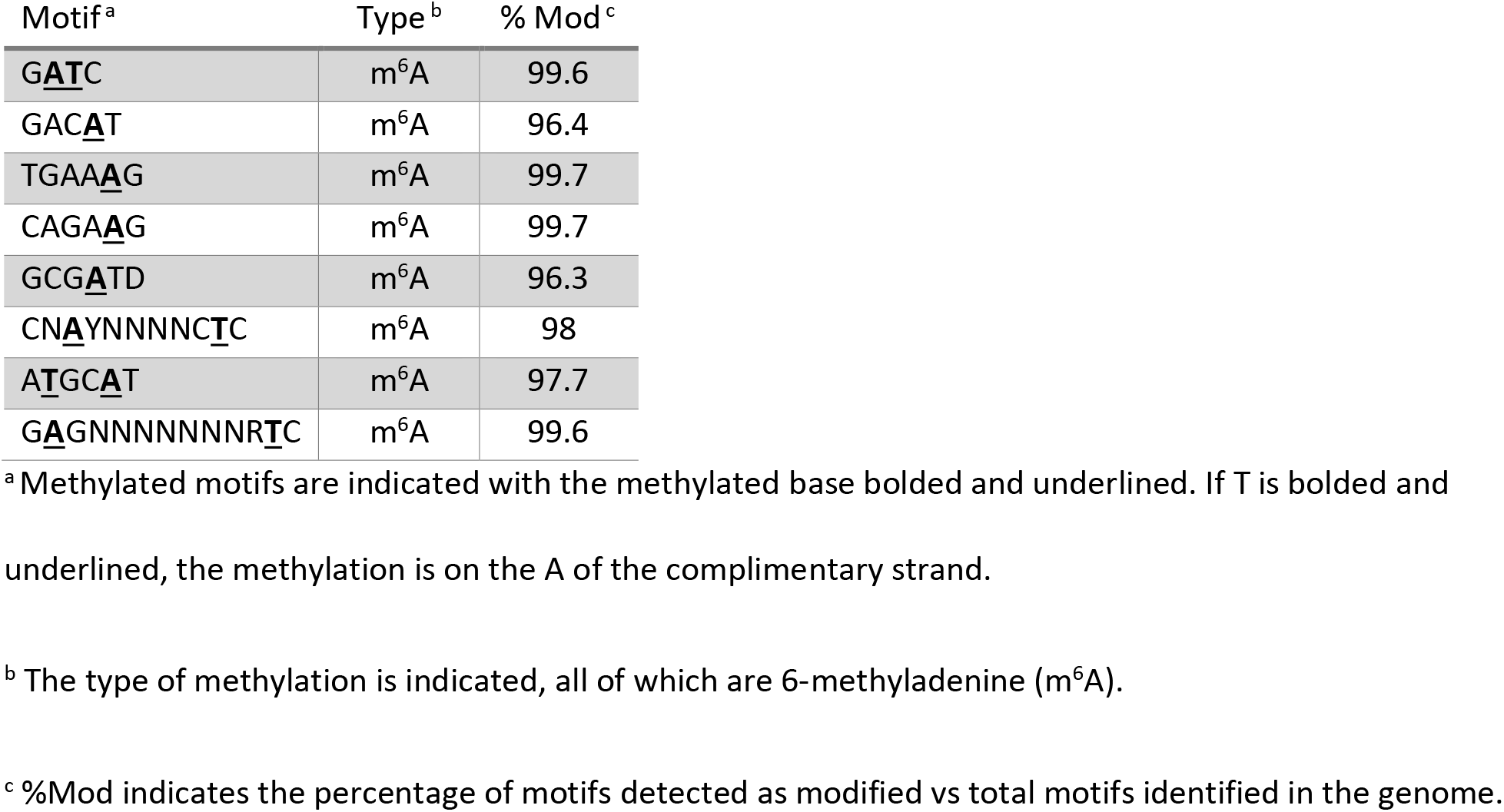
Methylome analysis of *C. clariflavum* 4-2a

To engineer *E. coli* to mimic the methylome of *C. clariflavum* to evade restriction, each identified methyltransferase was expressed from an arabinose-inducible promoter and inserted sequentially into the *E. coli* chromosome into different *attB* sites. The methyltransferases were cloned into the non-replicating integration vector with a single *attP* site. The methyltransferases were sequentially integrated into an *E. coli* strain that natively encodes Dam methyltransferase to methylate G(m^6^A)TC but lacks *dcm* because C(m^5^C)WGG is not methylated by *C. clariflavum*, resulting in strain AG5645 (Supplemental table 2). The plasmid backbone of each integrated vector was removed as described above using ΦC31 Int, allowing repeated use of the kanamycin resistance gene. Methylome analysis was performed to examine the functionality of each methyltransferase in *E. coli* (Table 2). Six motifs were identified; of these, four showed complete methylation, and two showed partial methylation. No methylation was detected for ATGC**A**T and CAGA**A**G, suggesting that some methyltransferases were not function in *E. coli*.

**Table 2:**
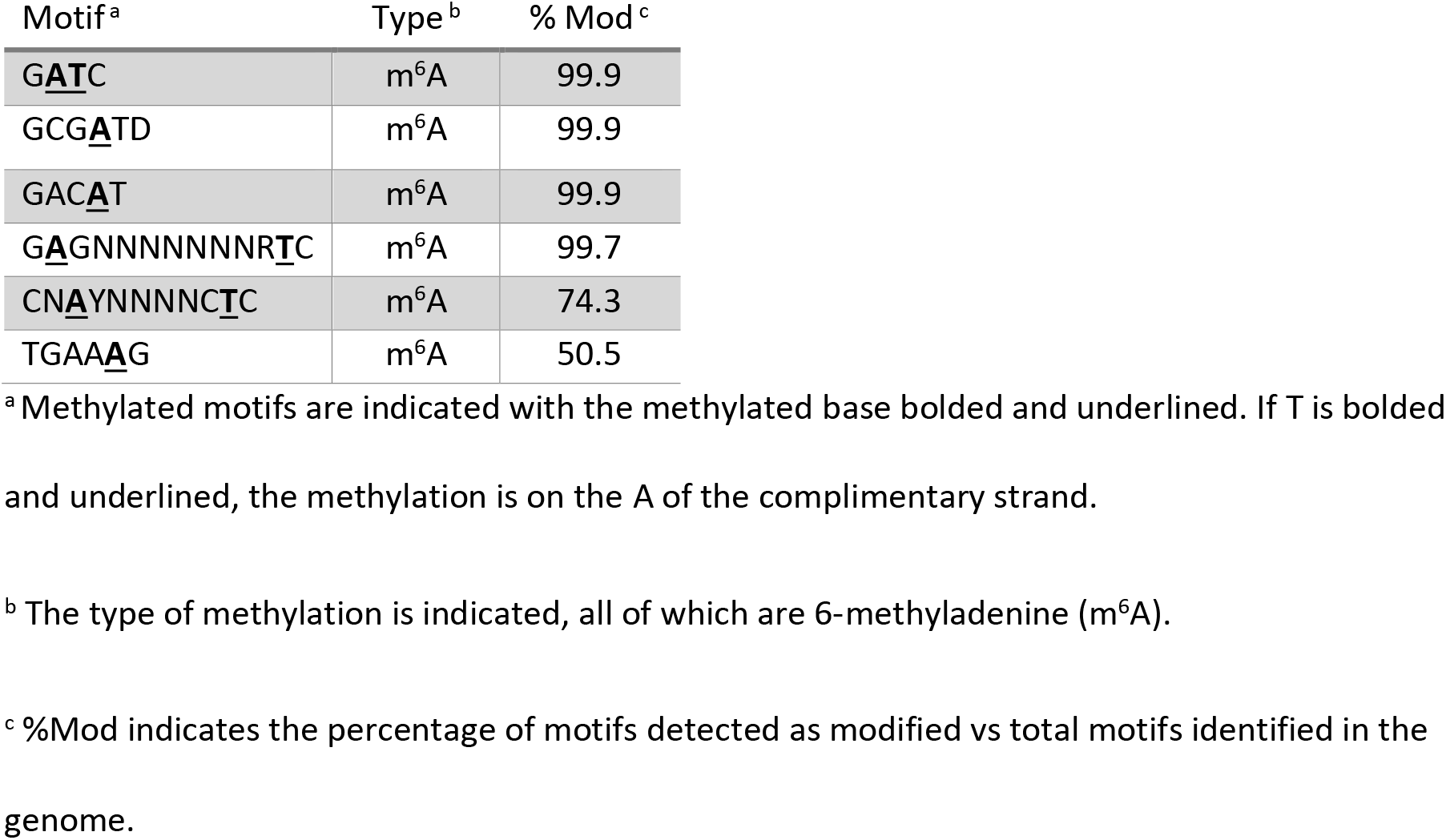
Methylome analysis of *E. coli* AG5645 strain to mimic methylation in *C. clariflavum* 4-2a

### Transformation of C. clariflavum 4-2a

Even though methylation in *E. coli* was incomplete, we attempted to transform *C. clariflavum* with plasmid DNA methylated by *E. coli* strain AG5645. To determine the impact of DNA methylation on transformation of *C. clariflavum*, plasmid pMTL83151 was isolated out of arabinose-induced *E. coli* methylation strain AG5477 and used to transform *C. clariflavum*. As a control, unmethylated plasmid was isolated out of Top10 *Δdcm::frt* and used to transform the strain. Unmethylated plasmid never yielded colonies, while the methylated plasmid yielded 9 ± 3 colony forming units per microgram of DNA (CFU/μg).

## DISCUSSION

The DNA integration system developed here expands the *E. coli* genetic toolbox to provide a quick and simple method to integrate up to twelve heterologous DNA constructs stably and sequentially into the chromosome of *E. coli*. In combination with the CRIM system, this enables insertion of up to sixteen plasmids. Integration by our system can occur via a single co-transformation of two plasmids, including one recombinase-expression plasmid that is common for all insertions into that *attB* site and one that can be customized to carry the desired cargo. Site-specific recombination inserts DNA into the chromosome more quickly than classic techniques like homologous recombination. Furthermore, an additional site-specific recombination step can be utilized to remove vector backbones and resistance markers, which avoids spontaneous resistant mutants than can hinder *sacB* and other counter-selection systems. Using this system enables stable chromosomal integration in one day and unmarked insertions in three days, while tools such as homologous recombination often requires several steps and at least five days.

Another characteristic of the system is the ability of the recombinases to recombine DNA efficiently and reliably. They function without a significant decrease in efficiency as the size of the insert increases; in our hands, cargo sizes upwards of 7 kb have been integrated, and no decrease in recombination efficiency was seen (unpublished data). Size-based limitations on DNA insertion should be primarily dependent on the diminishing efficiency of DNA entry into the cell as the DNA size increases. This overcomes some of the size-associated challenges with other genetic tools, like use of the lambda Red recombinase system [3] for heterologous DNA insertion. The ability to rapidly integrate constructs, at single copy, will help to increase the speed of engineering in *E. coli*, especially for high throughput, single copy library evaluation.

The DNA integration system described here contains other useful features. Like the CRIM system [8], it uses the conditional *oriR6K* origin of replication, which is only able to replicate in *E. coli* strains engineered to express the *pir* gene. This enables the integration vectors to act as a suicide vector in most *E. coli* strains, including the one containing the poly-*attB* cassette. Next, the vector backbone can easily be removed after DNA integration because the *oriR6K* origin and the antibiotic resistance marker are flanked by ΦC31*att* sites. Removal of the backbone allows for reuse of the antibiotic resistance marker and reduces repetitive DNA, diminishing the likelihood of homologous recombination events when stacking multiple genes into the poly-*attB* cassette. Another feature of this system is the unidirectional nature of the serine recombinases. Integration creates new attachment sites, *attL* and *attR*, and these new *att* sites are not recognized by the recombinases. This is especially beneficial when integrating multiple constructs in close proximity on the genome. Unlike the Flp-*frt* and Cre/*lox* systems, where the resulting scars are still substrates for the recombinase, ΦC31 does not act on the remaining ΦC31 *attR* sites after the marker is removed. Because these resulting *attR* scars cannot recombine with each other, there is no genome instability if two scars are near each other.

Initially, the first set of recombinase helper plasmids was constructed that allowed for integration in two steps. Using this approach, twelve of the recombinases were functional, but two were not. For unknown reasons, the BT and RV recombinases did not function as expected; this could be further explored in the future to increase the number of available sites for DNA insertion. To decrease the number of steps in this process, the ability to transform both plasmids in one event was desirable. The second set of helper plasmids enabled one-step integration, though two of the recombinases were not successfully cloned. We speculate that high expression of the Wβ and Spβ recombinases was toxic for *E. coli*. However, the availability of the first set of plasmids still allows the use of the Wβ and Spβ recombinases via the two-step process. Of the recombinases that are functional in the one-step process, R4, BxB1, and TG1 are highly efficient and reliable, so these three insertion sites should be targeted first when using this integration system. The ΦC1 and MR11 recombinases are least reliable, commonly integrating into pseudo-*att* sites and with lower efficiencies; therefore, these should be the last two sites to be considered. There is a decrease in transformation efficiency when co-transforming with the strong promoter helper plasmids, but the speed of the one-step process makes them more useful when speed is desired. The two-step approach will be most useful when high integration efficiency is needed, such as for the creation of large libraries.

Technologies to transform diverse non-model microorganisms are desperately needed, and mimicking the methylation patterns of the target organism to avoid restriction is an important approach to enable transformation. To this end, and as a demonstration of the utility of this tool, we engineered *E. coli* to express *C. clariflavum* methyltransferases to protect plasmid DNA and enable the genetic transformation of *C. clariflavum*. Transformation efficiency in this strain is still low, and further optimization will likely increase the efficiency. Two of the methylated motifs found in *C. clariflavum* were not found in the *E. coli* methylation strain, and two other motifs were methylated less than 100% in the *E. coli* genome, which likely impacts the transformation efficiency. Because *C. clariflavum* is a thermophilic organism, the activity of the methyltransferases at *E. coli* growth temperatures could decrease functionality. Future utilization of mesophilic methyltransferases that target the same motifs or engineered versions of the thermophilic enzymes to increase activity at mesophilic temperatures could help to overcome this challenge. Even with the incomplete methylation, low transformation efficiency was enabled, which allows for further genetic manipulation of *C. clariflavum* for fundamental studies and metabolic engineering related to lignocellulose deconstruction and bioconversion.

Finally, the tools developed here are directly applicable to other organisms, including non-model microbes. Because these recombinases do not require any host factors to function, this system should be adaptable to any strain in which the poly-*attB* “landing pad” can be inserted into the chromosome. Recently, this has been developed for various Gamma- and Alpha-proteobacteria in a toolset called Serine recombinase-Assisted Genome Engineering (SAGE) [33]. Together, these toolsets will help accelerate strain engineering across diverse organisms.

## Acknowledgements

Funding was provided in part by The BioEnergy Science (BESC) and The Center for Bioenergy Innovation (CBI), U.S. Department of Energy Bioenergy Research Centers supported by the Office of Biological and Environmental Research in the DOE Office of Science. Funding was also provided in part by Office of Biological and Environmental Research in the DOE Office of Science under Award Number DE-SC0019401. Oak Ridge National Laboratory is managed by UT-Battelle, LLC, for the U.S. DOE under contract DE-AC05-00OR22725. The funders had no role in study design, data collection and analysis, decision to publish, or preparation of the manuscript.

---

**Supplemental Table S1:** All strains used with their strain collection number, genotype, and description.

| Strain # | Genotype | Description |
| --- | --- | --- |
| ***E. coli* GT115** | *F- mcrA Δ(mrr-hsdRMS-mcrBC) φ80lacZΔM15 ΔlacX74 nupG recA1 araD139 Δ(ara-leu)7697 galE15 galK16 rpsL(StrA) endA1 Δdcm uidA(ΔMluI)::pir-116 ΔsbcC-sbcD* | *pir+* stain for propagation of plasmids containing oriR6K |
| **AG583** | *E. coli* Top10 Δdcm::f<sub>rt</sub> |  |
| **AG2004** | *E. coli* GT115 + pLAR080 | pAH144+poly-*attB* (*aad+*) |
| **AG2005** | *E. coli* AG 583 HK::poly-*attB* | Poly-*attB* in the chromosome of an *araC- dcm-* strain |
| **AG3525** | *E. coli* WM3188 Δdcm::f<sub>rt</sub> HK::poly-*attB* | Poly-*attB* in the chromosome of an *araC+ dcm-* strain |
| **AG4277** | *E. coli* BW25113 Δ*mcrA*::f<sub>rt</sub> Δ*mcrC-mrr*::f<sub>rt</sub> Δdcm::f<sub>rt</sub> Δ*dam*::f<sub>rt</sub> HK::poly-*attB* | Poly-*attB* in the chromosome of an *araC+ dam- dcm-* strain |
| **AG1434** | *E. coli* GT115 + pGSs009 | Non-replicating integration plasmid with the P<sub>T5lac</sub> and lacZα |
| **AG3409** | *E. coli* GT115 + pLAR067 | Non-replicating integration plasmid with P<sub>BAD</sub> and lacZα |
| **AG3273** | *E. coli* GT115 + pLAR031 | Non-replicating *attP*-cassette plasmid |
| **AG5425** | *E. coli* GT115+ pLAR140 | pGSs009 with BxB1 *attP* site |
| **AG1460** | *E. coli* AG583 + pGSs037 | pINT-TS ΦC31 |
| **AG1461** | *E. coli* AG583 + pGSs038 | pINT-TS RV |
| **AG1463** | *E. coli* AG583 + pGSs040 | pINT-TS TG1 |
| **AG1464** | *E. coli* AG583 + pGSs041 | pINT-TS BxB1 |
| **AG1484** | *E. coli* AG583 + pGSs042 | pINT-TS A118 |
| **AG1485** | *E. coli* AG583 + pGSs043 | pINT-TS ΦC1 |
| **AG1486** | *E. coli* AG583 + pGSs044 | pINT-TS ΦK38 |
| **AG1487** | AG583 + pGSs045 | pINT-TS MR11 |
| **AG1488** | AG583 + pGSs046 | pINT-TS R4 |
| **AG1489** | AG583 + pGSs047 | pINT-TS SPβc |
| **AG1490** | AG583 + pGSs048 | pINT-TS TP901-1 |
| **AG1492** | AG583 + pGSs050 | pINT-TS Φ370 |
| **AG1495** | AG583 + pGSs053 | pINT-TS BL3 |
| **AG1496** | AG583 + pGSs054 | pINT-TS ΦBT1 |
| **AG1524** | AG583 + pGSs082 | pINT-TS Wβ |
| **AG3463** | AG583 + pLAR074 | p<sub>Eno</sub>-BxB1 |
| **AG3512** | AG583 + pLAR052 | p<sub>Eno</sub>-Φ370 |
| **AG3513** | AG583 + pLAR053 | p<sub>Eno</sub>-A118 |
| **AG3515** | AG583 + pLAR055 | p<sub>Eno</sub>-ΦC1 |
| **AG3516** | AG583 + pLAR056 | p<sub>Eno</sub>-ΦK38 |
| **AG3517** | AG583 + pLAR057 | p<sub>Eno</sub>-MR11 |
| **AG3518** | AG583 + pLAR058 | p<sub>Eno</sub>-R4 |
| **AG3520** | AG583 + pLAR060 | p<sub>Eno</sub>-TG1 |
| **AG3521** | AG583 + pLAR061 | p<sub>Eno</sub>-TP901-1 |
| **AG3522** | AG583 + pLAR062 | p<sub>Eno</sub>-BL3 |
| **AG3524** | AG583 + pLAR047 | p<sub>Eno</sub>-ΦC31 |
| **AG 5407** | GT115 + pMTV37 | pLAR067 + CloclaDRAFT _2133-R4 |
| **AG5408** | GT115 + pMTV38 | pLAR067 + CloclaDRAFT _2368-2369-phiK38 |
| **AG5409** | GT115 + pMTV39 | pLAR067 + CloclaDRAFT cla_1830-1831-BXB1 |
| **AG5410** | GT115 + pMTV40 | pLAR067 + CloclaDRAFT _1058-TP901 |
| **AG5411** | GT115 + pMTV41 | pLAR067 + CloclaDRAFT _1868-1869-TG1 |
| **AG5412** | GT115 + pMTV42 | pLAR067 + CloclaDRAFT _1961-BL3 |
| **AG5586** | GT115 + pMTV43 | pLAR067 + CloclaDRAFT _1058-A118 |
| **AG5456** | GT115 + pMTV44 | pLAR067 + CloclaDRAFT _1994-phi370 |
| **AG5645** | BW25113 Δ*mcrA*::*frt* Δ*mcrC-mrr*::*frt*<br>Δ*dam*::*frt* Δ*dcm*::*frt* HK::pLAR080<br>R4::CloclaDRAFT_2133 Φk38:: CloclaDRAFT<br>_2368-2369 BXB1:: CloclaDRAFT _1830-1831<br>TP901-1:: CloclaDRAFT _1058 TG1::<br>CloclaDRAFT _1868-1869 BL3:: CloclaDRAFT<br>_1961 Φ370:: CloclaDRAFT _1994 A118::<br>CloclaDRAFT _1058 | *E. coli* strain expressing *C. clariflavum*<br>4-2a methyltransferases |
| ***Clostridium clariflavum***<br>**4-2a (AG1933)** | Wild type | [24] |

**Supplemental Table S2:**
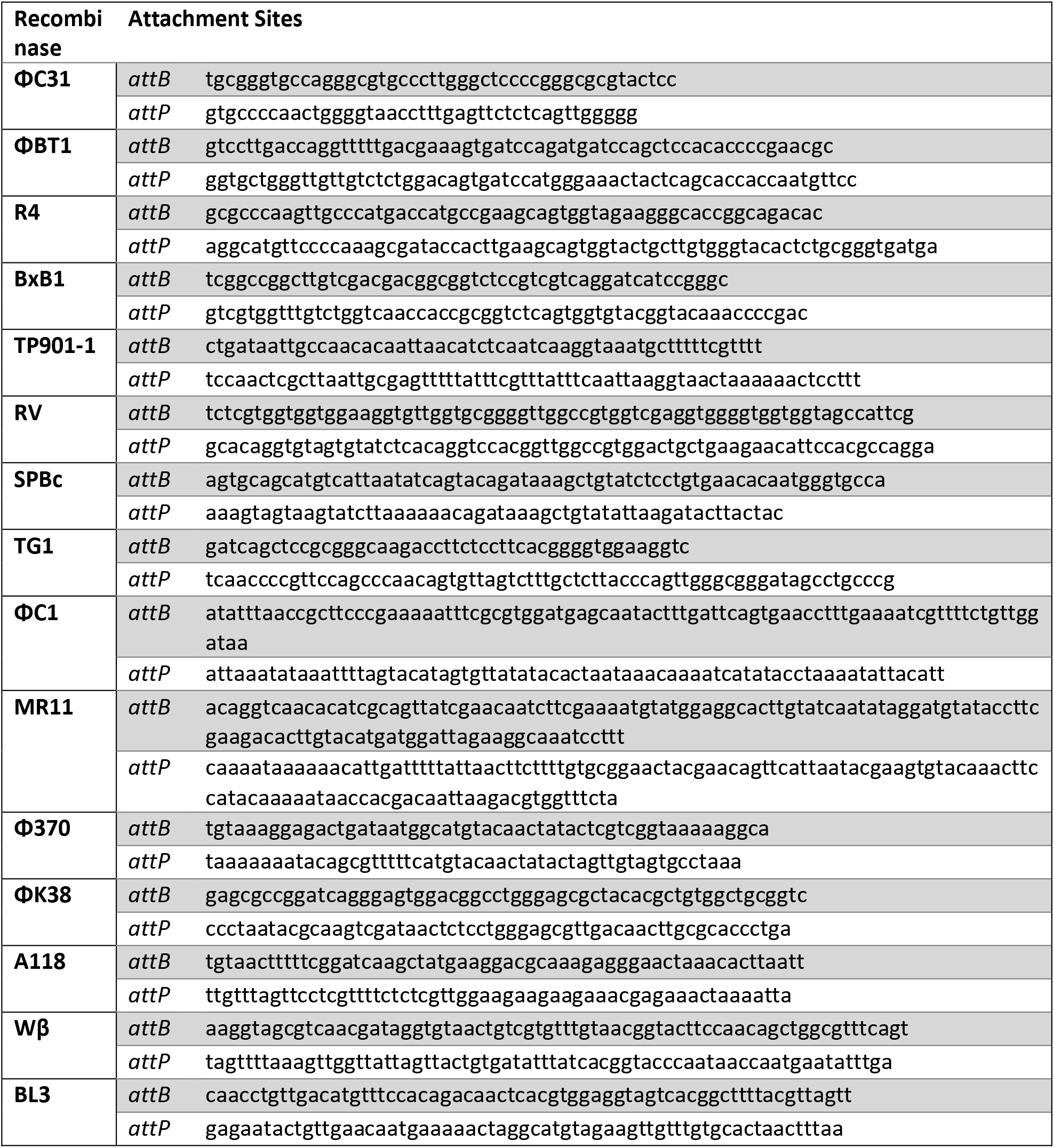
Attachment sites for each recombinase used. Previously described[7]

**Supplemental Table S3:**
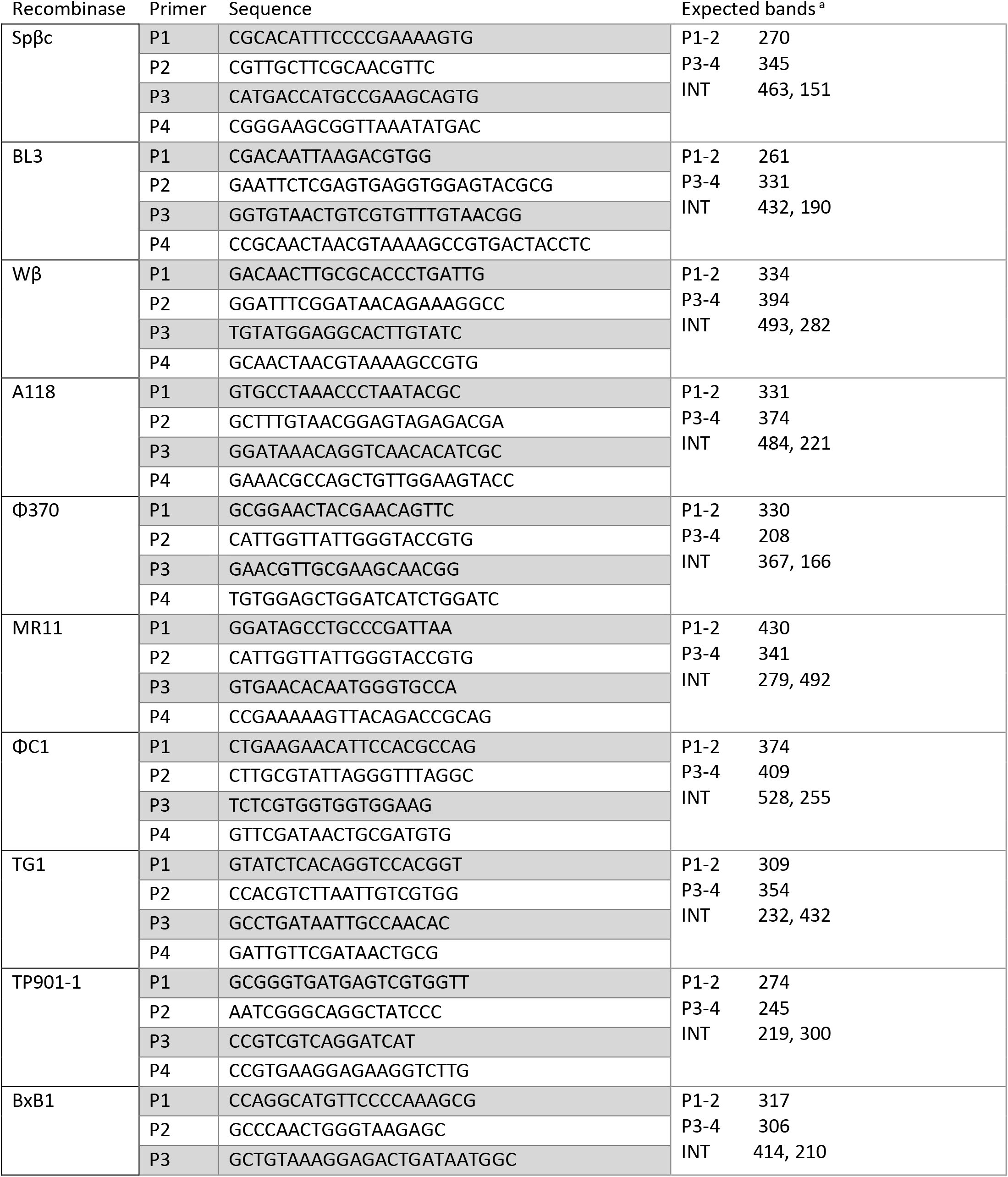
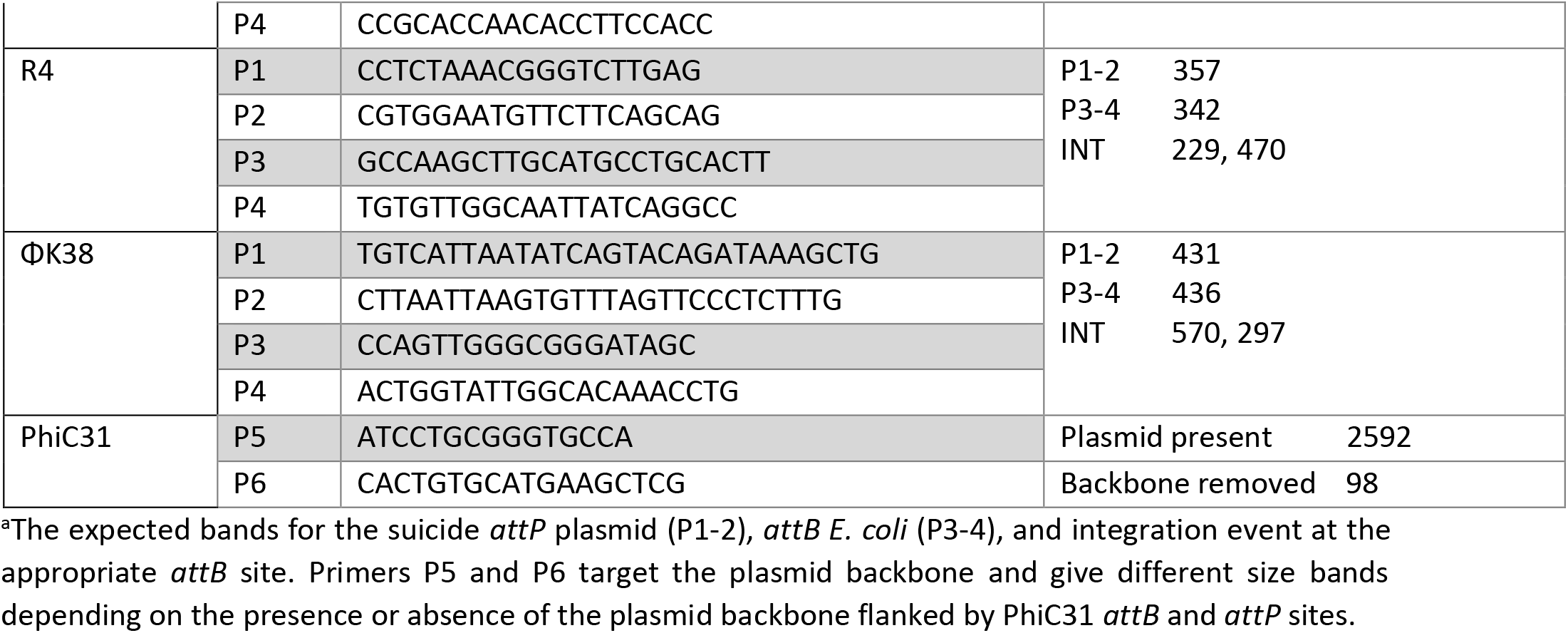
Primers used to determine integration into each *attB* site.

**Supplementary Figure 1.**
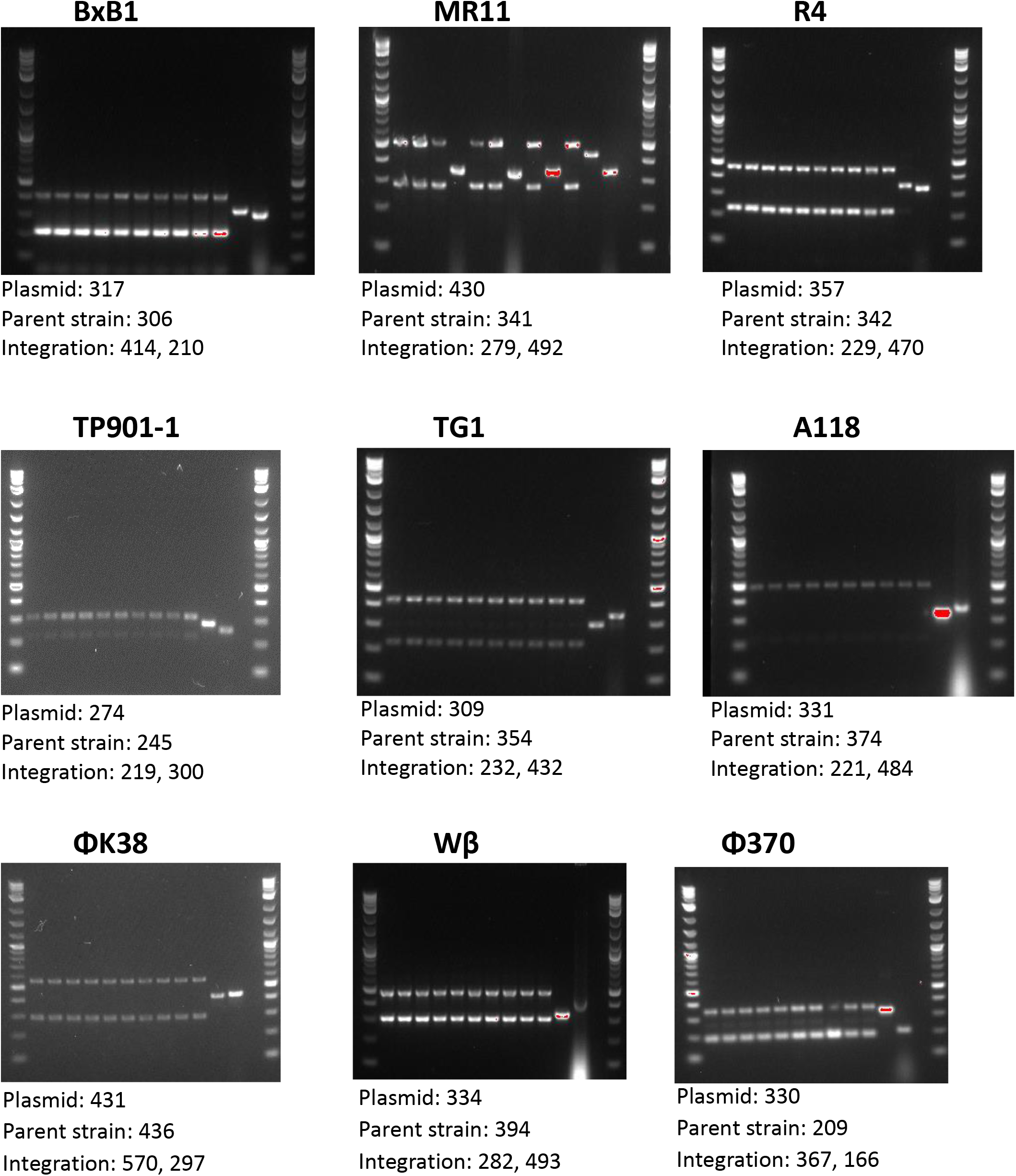
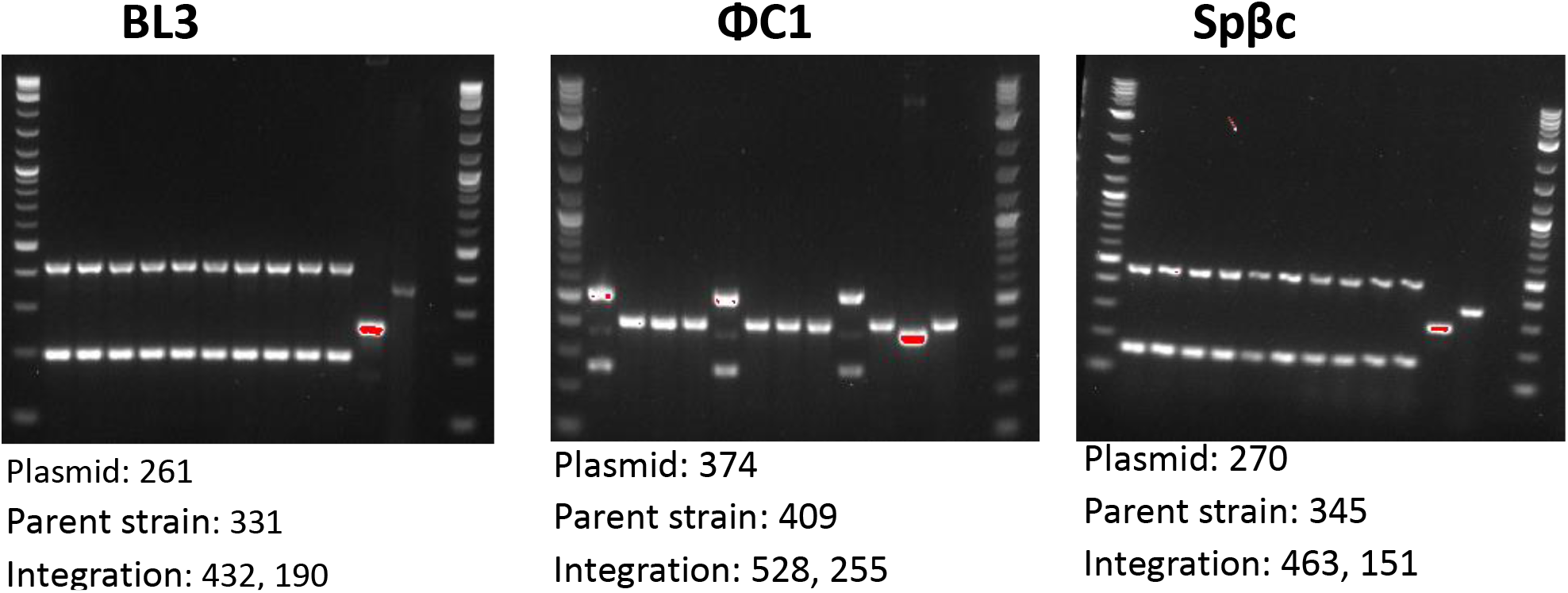
PCR assay to determine if integration occurred at the correct locus. The recombinase being tested is listed above each agarose gel image. For each gel, Lane 1: NEB 1kb plus ladder, Lanes 2-11: transformed colonies being screened for correct insertion, Lane 12: pLAR031 plasmid control (intact *attP* site), and Lane 13: Parent strain control (intact *attB* site). Under each gel image, the expected band sizes are listed (in base pairs) for the plasmid control (having an intact *attP* site), parent strain control (intact *attB* site), and for successful integration.

**Supplementary Figure 2.**
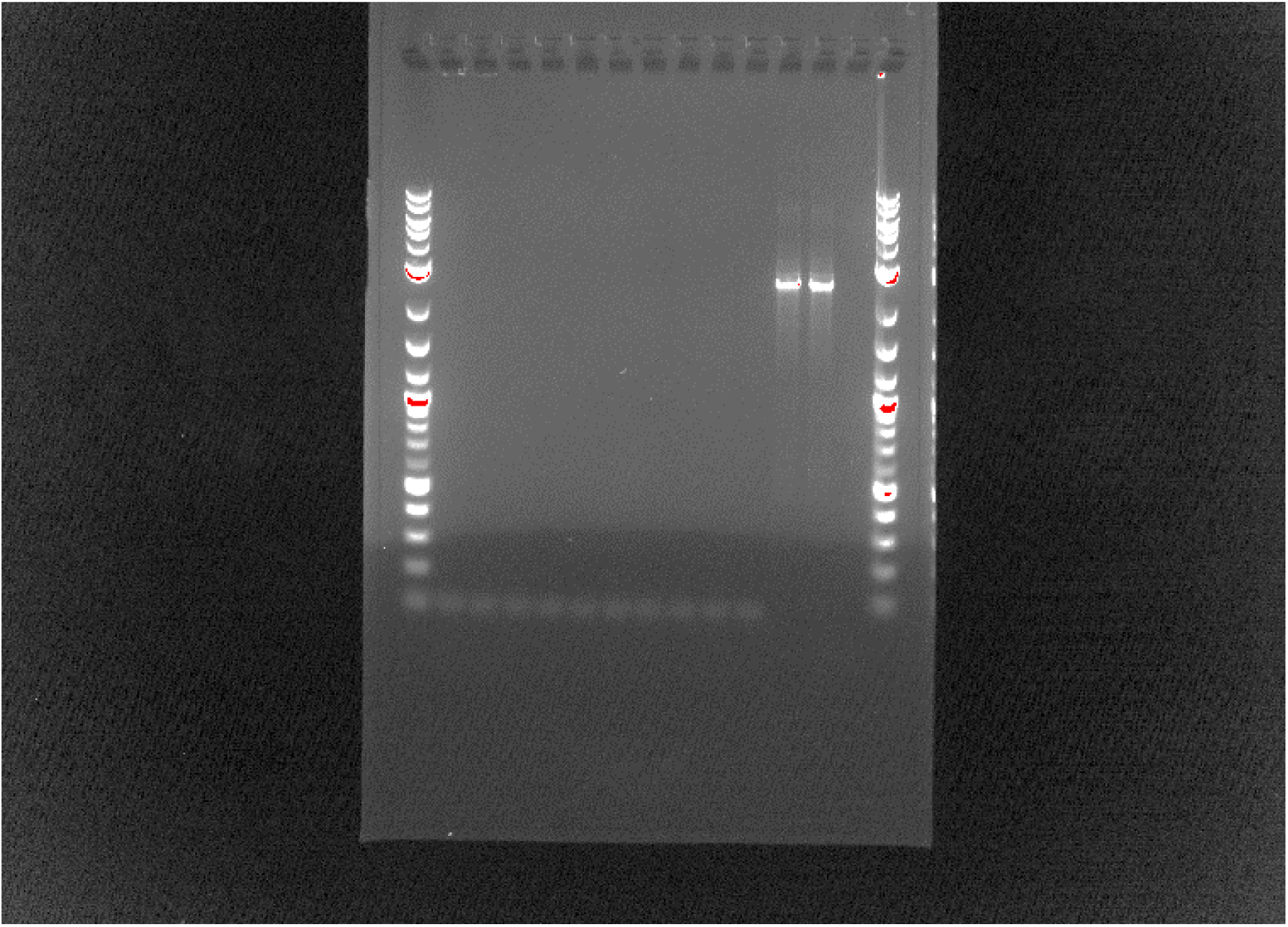
PCR assay to determine if transient ΦC31 Int expression resulted in loop-out of the cargo plasmid backbone. Lane 1: NEB 1kb plus ladder, Lanes 2-11: transformed colonies being screened for loss of ΦC31 *attB-kanR-attP* cassette, Lane 12: pLAR031 plasmid control (with intact ΦC31 *attB-kanR-attP cassette*), and Lane 13: Parent strain control (strain AG2005 R4*att*::pLAR031, which has intact ΦC31*attB-kanR-attP* cassette), Lane 14: blank, Lane 15: ladder. Expected band sizes are (in base pairs) for the plasmid control and parent strain control (with intact ΦC31 *attB-kanR-attP cassette*), 2592 bp; excised plasmids backbone, 98 bp.

